# A Novel Mycovirus Identified in Clinical Isolates of *Cryptococcus neoformans*

**DOI:** 10.64898/2026.08.24.746608

**Authors:** Mysha N. Turk, Abberly E. Dela Rosa, J. T. Graham Solomons, Virginia E. Glazier

## Abstract

Mycoviruses are widespread throughout the fungal kingdom and are known to infect diverse fungal taxa including fungal species that are important plant and human pathogens. Although many mycoviruses have been found to have minimal effects on their host, several viruses have been found to modulate fungal physiology, and as a result impact fungal virulence. Screens for mycoviruses in clinically relevant fungi have identified numerous mycoviruses within several important human pathogens, however mycoviruses remain uncharacterized in the clinically relevant human pathogen *Cryptococcus neoformans. C. neoformans* is an opportunistic encapsulated yeast responsible for life-threatening cryptococcal meningitis, a leading cause of mortality among immunocompromised individuals, particularly those with HIV/AIDS. We performed a search for viral RNA-dependent RNA Polymerase (RdRP) signatures in publicly available *C. neoformans* transcriptomic data. This search identified *Totiviridae* viral genomes within six clinical isolates of *C. neoformans* from Botswana. All six isolates originated from the CSF of HIV positive individuals with cryptococcal meningitis. Reverse transcription PCR (RT-PCR) independently validated the continued presence of the virus in three of these clinical isolates. Subsequent analysis of the viral genome identified two genotypes of a single species of Totivirus. This new species possesses canonical features of the *Totiviridae* family, including a slippery heptamer and a predicted RNA pseudoknot structure involved in programmed −1 ribosomal frameshifting for RdRP expression. Taken together, these results provide evidence of a mycovirus capable of infecting *C. neoformans*.

**Importance:** We report the identification of a mycovirus capable of infecting *Cryptococcus neoformans*. While mycoviruses inhabit diverse fungal taxa, they have previously remained uncharacterized in this clinically significant fungal pathogen. The identification of the mycoviruses within *C. neoformans* isolated from the CSF of individuals with cryptococcal meningitis places emphasis on the potential role of mycoviruses in the pathogenesis of *C. neoformans*.

## Observation

*Cryptococcus neoformans* is a clinically relevant fungal pathogen that can cause cryptococcal meningoencephalitis in immunocompromised individuals, such as those with HIV/AIDS. Given the high degree of variability observed in cryptococcal meningitis disease presentation, development of drug resistance, and patient outcomes, studies have increasingly focused on the differences between clinical isolates of *C. neoformans* as influencing infection outcome (1–4). A missing component in understanding clinical isolate diversity and its impact on disease outcome is the potential presence of mycoviruses within *C. neoformans* isolates.

Mycoviruses encompass a broad range of viral families and fungal hosts. The impact of mycoviruses on their fungal host is varied and ranges from latent infection to dramatic phenotypic changes that induce either hypervirulence or hypovirulence depending on the specific viral-host pairing (5). The explosion of publicly available sequencing data has emerged as a powerful resource for identifying mycoviruses in clinical and environmental isolates, facilitating the discovery of novel mycoviruses across diverse fungal lineages (6, 7).

To identify potential mycoviruses in *C. neoformans*, we screened the publicly available NCBI Sequence Read Archive (SRA) data for transcriptomic studies in the Serratus Explorer database which aligns SRA libraries to *rdp1* using DIAMOND (8). The Serratus database contains 2940 *Cryptococcus neoformans* transcriptomic SRA files. Our screening identified six clinical isolates that contained RNA Dependent RNA Polymerase (RdRP) signatures (Fig. 1A). All six clinical isolates were retrieved from the CSF of HIV+ individuals with cryptococcal meningitis seeking treatment at hospitals in Gaborone or Francistown Botswana (2). All belong to serotype A with both A and alpha mating types present (9). The clinical isolates were obtained from a study that characterized the transcriptomic profiles of the strains under various conditions including directly from the patient CSF (hCSF) and grown using artificial CSF (aCSF), rabbit CSF (rCSF), YPD or Capsule-inducing medium (CAP). Interestingly, some of the isolates tested positive for the presence of RdRP signatures under multiple transcriptomic profile conditions, NRHc5028 (6 positive samples), NRHc5010 (4 positive samples), while others only contained RdRP signatures under one condition (NRHc5027, NRHc5030, PMHc1033 and PMHc1063).

**FIG 1.**
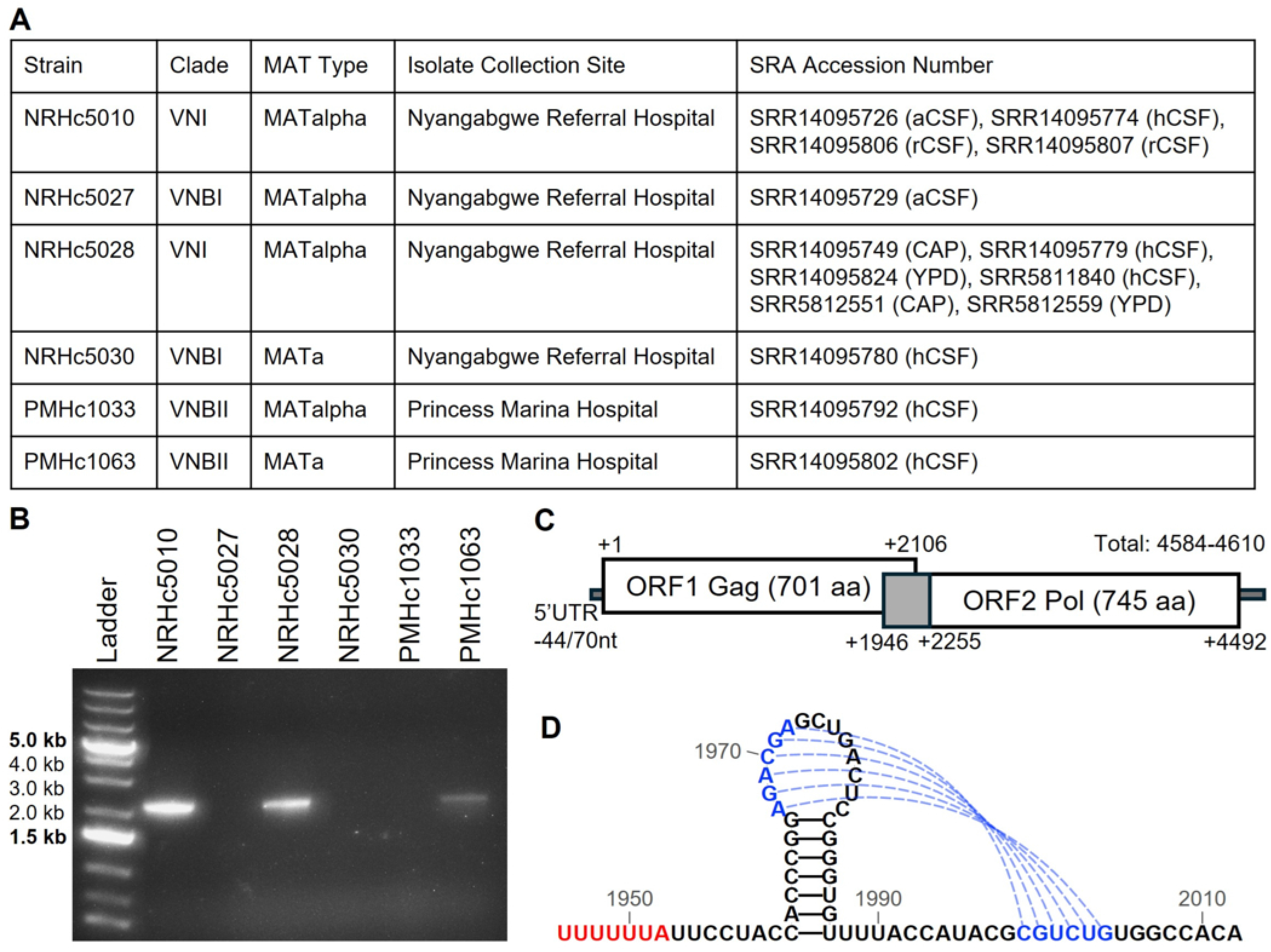
Identification of a mycovirus in clinical isolates of *C. neoformans* (A) Table containing strain information for RdRP positive *C. neoformans* SRA libraries (B) Agarose gel electrophoresis (1% agarose, 1× TAE) of RT-PCR amplicons targeting the ORF2-encoded RNA-dependent RNA polymerase (RdRP) region. Bands correspond to the expected size of the targeted viral RdRP fragment (2121bp). (C) Schematic of the dsRNA genome showing ORF1 (Gag, capsid protein) and ORF2 (Pol, RdRP), (D) The predicted secondary structure of the −1 ribosomal frameshift region made in RNAcanvas.

To validate the *in silico* findings, we obtained the six *C. neoformans* isolates from Dr. John Perfect’s laboratory and evaluated them *in vitro*. Using RT-PCR amplification of cDNA targeting the ORF2-encoded RdRP region followed by Sanger sequencing, we successfully confirmed the presence of the virus in NRHc5010, NRHc5028, and PMHc1063 (Fig. 1B). Isolates NRHc5027, NRHc5030, and PMHc1033 failed to have the ORF2-encoded RdRP amplified via RT-PCR. The failure to amplify viral genomic material from these isolates aligns with the observation that their specific RdRP signatures were restricted to a single transcriptomic profiling sample. These results taken together may suggest either a lower limit of detection of the viral genome due to low copy number, or the loss of the virus in some or all of the cells of a given isolate during isolate maintenance and passaging. This potential variation in viral retention and copy number across clinical isolates may be the result of isolate variability in *C. neoformans* host defense mechanisms. While *Cryptococcus neoformans* natively possesses a functional RNA interference (RNAi) system used for antiviral defense, recent work has demonstrated that spontaneous loss of RNAi proficiency is a common evolutionary trajectory among clinical isolates (4). This loss of RNAi functionality would presumably increase susceptibility to viral infection. Remarkably, the clinical isolate identified in our study with the highest retention of viral RdRP signatures, NRHc5028, was found by Huang *et al*. to lack the siRNAs required for functional RNAi silencing. We therefore postulate that the presence or absence of functional RNAi machinery within these specific clinical isolates could provide a mechanistic explanation for the observed isolate variability in viral genome retention.

Assembly of the viral genomes from the RNA sequencing data using SPAdes v3.15.3 revealed that the six isolates form two genotypes with Genotype A identified in isolates NRHc5010, NRHc5027, NRHc5030, PMHc1033, and PMHc1063 and Genotype B found in isolate NRHc5028. CLUSTALW was used to align the viral genomes, with the variation between the two strains being 145 SNPs, resulting in approximately 96% sequence similarity between the two genotypes. Minor heterogeneity was observed in the 5′ UTR terminus, where Genotype B (isolate NRHc5028) possessed an additional 26 nucleotides. We are therefore classifying the two genotypes as two distinct strains of the same novel viral species. NCBI ORF Finder was used to predict putative open reading frames (ORFs). The viral genomes contained two ORFs that matched the known Totivirus capsid proteins (ORF1 Gag) and RdRP (ORF2 Pol) encoding genes using BLAST. Further analysis of the viral genome revealed characteristics found in other members of the Totivirus family including Histidine 160 of the capsid protein which is a conserved residue required for cap snatching (10). Evidence of structural characteristics required for a −1 ribosomal frameshift can be observed in the presence of the slippery heptamer UUUUUUA, a seven nucleotide spacer followed by a hairpin that forms an H-type pseudoknot (Fig. 1C) (11, 12).

The viral sequence exhibits high homology within the family *Totiviridae* which was confirmed by both Serratus explorer and localized BLAST of ORF1 and ORF2. To precisely establish its evolutionary relationships and narrow down its taxonomic placement across the family *Totiviridae*, a comprehensive phylogenetic analysis was performed based on the conserved RdRP region. Broad alignments were iteratively sub-sampled to eliminate redundant sequences and further narrow down the reference dataset of representative totiviruses in the *Totivirus* genus (Fig. 2). The analysis resolved four well-supported clades corresponding to the previously established subgroups within the *Totivirus* genus (13). The RdRP sequences analyzed in this study are nested within subclade I-B. Phylogenetic relationships indicates that this mycovirus is most closely related to other characterized totiviruses that infect fungal hosts, including *Malassezia sympodialis mycovirus 1, Erysiphe necator-associated totivirus 8*, and *Scheffersomyces segobiensis virus L*. Based on its definitive placement within the Totivirus genus, its novel host, and its distinct genetic profile, we propose the name *Cryptococcus neoformans totivirus 1* (CnTV1) for this novel viral species.

**FIG 2.**
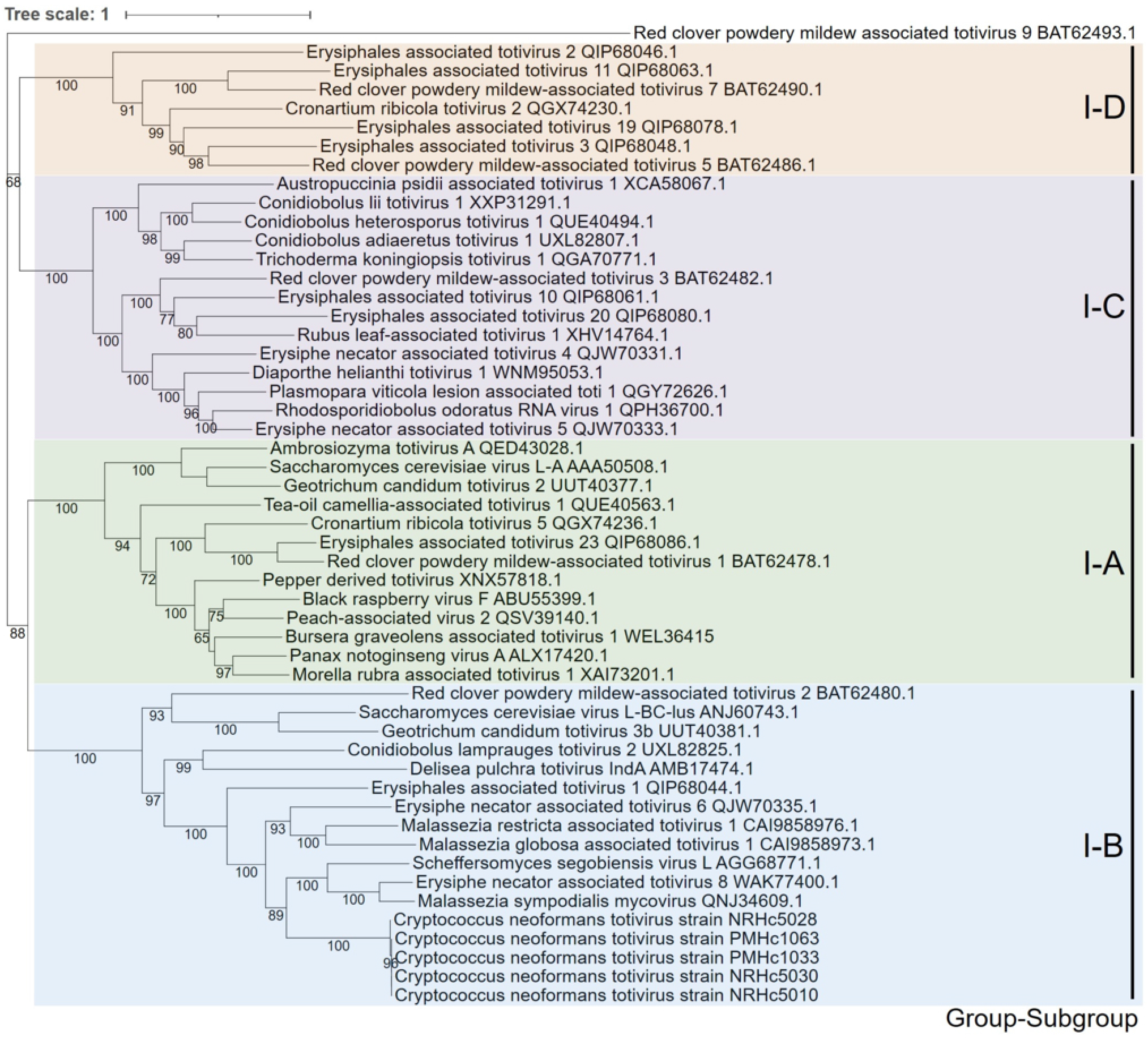
Phylogenetic relationship of *Cryptococcus neoformans totivirus 1* isolates within the *Totivirus* genus. The maximum-likelihood tree was constructed using representative RNA-dependent RNA polymerase (RdRP) amino acid sequences. The four major established subgroups within the genus *Totivirus* group I are indicated, with *Red clover powdery mildew associated totivirus 9* forming the outgroup and representing group II. Node labels represent bootstrap support values from 1,000 replicates. Scale bar indicates amino acid substitutions per site.

This study showcases the wealth of uncharacterized biological information hidden within public data repositories like the Sequence Read Archive (SRA). Leveraging these pre-existing datasets provides a rapid and highly accessible pathway for viral discovery (6, 7, 14). Applying this pipeline to clinical isolates of *Cryptococcus neoformans* led to the discovery of CnTV1, highlighting the value of sampling diverse strains. Characterizing patient-derived isolates in particular has immense utility, as it may reveal the underlying biological factors driving the high variability in disease presentations and outcomes observed in the clinic (1, 2). Understanding this variability requires investigating evolutionary mechanisms that generate phenotypic diversity among strains. Along these lines, given that a decrease or loss of RNA interference (RNAi) proficiency among clinical isolates of *C. neoformans* can drive the mobilization of transposable elements, thereby facilitating rapid drug resistance while simultaneously increasing susceptibility to viral infections, the interplay between host RNAi defense and mycovirus carriage represents a new dimension to our understanding of fungal pathogenesis (4). Ultimately, continuing to investigate isolate variability, such as the presence of viral elements, will be essential to a comprehensive understanding of *Cryptococcus neoformans* biology, virulence, and evolution.

### Fungal Culturing Conditions

Upon receipt of the *Cryptococcus neoformans* clinical isolates from Dr. John Perfect’s laboratory, cultures were initially cultured on Yeast Peptone Dextrose (YPD) agar plates then stored in YPD-Glycerol at −80°C. For molecular applications, clonal isolates were inoculated into liquid YPD medium and grown overnight at 30°C in a shaking incubator.

### RNA Isolation and RT-PCR Validation

Total RNA was extracted from overnight liquid cultures using TRIzol Reagent (Invitrogen). Next, 1,000 ng of total RNA was DNase-treated and reverse-transcribed into cDNA using the iScript cDNA Synthesis Kit (Bio-Rad). Targeted PCR amplification of the ORF2-encoded RdRP region with forward primer 5’-ACTCCTCCATCTCCCCTGCC-3’ and reverse primer 5’-AAAACCGAGCACGACAAACCCC-3’ was performed using 5 PRIME HotMaster Taq DNA Polymerase (Quantabio) denaturation 94°C for 2 min; followed by 40 cycles of denaturation at 94°C for 20 s, annealing at 54°C for 20 s, and extension at 68°C for 3 min with a final extension step of 68°C for 5 min. The resulting PCR products were resolved via electrophoresis on a 1% agarose TAE gel. Target bands were excised, purified using the QIAquick Gel Extraction Kit (Qiagen), and submitted for Sanger sequencing.

### Bioinformatics and Phylogenetic Reconstruction

Representative RNA-dependent RNA polymerase (RdRP) amino acid sequences were retrieved from the NCBI database and aligned with the RdRP from the assembled viral genomes. Sequence alignment was performed using MAFFT (v7.511) utilizing the L-INS-i settings (15). A maximum-likelihood phylogenetic tree was inferred using the IQ-TREE webserver under the LG+F+G4 evolutionary model, with branch support assessed using 1,000 bootstrap replicates and 1,000 SH-aLRT tests. The resulting phylogenetic tree was visualized and annotated using iTOL, with node labels indicating bootstrap support values.

## Data Availability

The assembled and annotated viral genomes for *Cryptococcus neoformans totivirus 1* Genotype A and Genotype B have been deposited in the NCBI Third-Party Annotation (TPA) database under accession numbers [TPA GENOTYPE A] and [TPA GENOTYPE B]. The consensus assemblies were derived from primary raw sequence data publicly available within the NCBI Sequence Read Archive (SRA) repository under BioProject accession PRJNA718309.

## Acknowledgements

We would like to thank Dr. John Perfect’s laboratory (Duke University School of Medicine) for generously providing the *Cryptococcus neoformans* clinical isolates used in the *in vitro* experiments.

